# Morphometric relationships and their contribution to biomass and cannabinoid yield in hybrids of hemp (*Cannabis sativa*)

**DOI:** 10.1101/2021.05.09.443329

**Authors:** Craig H. Carlson, George M. Stack, Yu Jiang, Bircan Taşkıran, Ali R. Cala, Jacob A. Toth, Glenn Philippe, Jocelyn K.C. Rose, Christine D. Smart, Lawrence B. Smart

## Abstract

The breeding of hybrid cultivars of hemp (*Cannabis sativa* L.) is not well described, especially the segregation and inheritance of traits that are important for yield. A total of 23 families were produced from genetically diverse parents to investigate the inheritance of morphological traits and their association with biomass accumulation and cannabinoid yield. In addition, a novel classification method for canopy architecture was developed. The strong linear relationship between wet and dry biomass provided an accurate estimate of final dry stripped floral biomass. Of all field and aerial measurements, basal stem diameter was determined to be the single best selection criterion for final dry stripped floral biomass yield. Along with stem diameter, canopy architecture and stem growth predictors described the majority of the explainable variation of biomass yield. Within-family variance for morphological and cannabinoid measurements reflected the heterozygosity of the parents. While selfed populations suffered from inbreeding depression, hybrid development in hemp will require at least one inbred parent to achieve uniform growth and biomass yield. Nevertheless, floral phenology remains a confounding factor in selection because of its underlying influence on biomass production highlighting the need to understand the genetic basis for flowering time in the breeding of uniform cultivars.

**Highlight:** Stem and canopy architecture traits are superior predictors of floral biomass yield and offer a good indication of hybrid uniformity in field plantings of genetically diverse cannabinoid hemp populations.

## Introduction

Hemp (*Cannabis sativa* L.) is a multipurpose crop with incipient potential in diverse markets. Hemp is a dioecious annual (2n=20) (Hirata, 1924) that along with *Humulus* spp., diverged from a common ancestor ca. 27 million years ago (McPartland, 2018). Generally thought to have been domesticated in Central Asia, its spatial distribution was reshaped by humans who have used it for millennia as a source of food, fiber, and medicine (Vavilov, 1926; Warf, 2014). The current scientific consensus is that the genus is monotypic (Small, 1972), however, niche populations do exist, primarily differentiated by latitude. What are likely remnants of early 20th century cultivation of fiber hemp, naturalized, locally-adapted populations can be found throughout the US. Nevertheless, collection and genetic characterization of wild populations has been extremely limited (Wenger *et al*., 2020), especially those in the native range (Soorni *et al*., 2017).

While hemp is dioecious with male heterogamety (XY) (Moliterni *et al*., 2004), instances of monoecy are commonplace. In an effort to increase grain yield, monoecious cultivars were first developed in European breeding programs. Monoecy is expressed in homogametic females (XX) but the ratio of staminate to pistillate flowers is quantitative and under autosomal control (XX+A) (Menzel, 1964). Presently, the genomic basis of sex determination (Petit *et al*., 2020) and monoecious expression (Faux *et al*., 2014) in *C. sativa* is not well understood. While monoecious cultivars are favored in grain production, all-female populations are preferred by growers of cannabinoid hemp because pollination can dramatically reduce cannabinoid yield (Small, 2015). Feminized populations are routinely produced by application of an ethylene inhibitor to one of the two female parents of a cross, which stimulates staminate flower formation, and all-female progeny that lack Y chromosomes. This simple technique, pioneered by Ram and Sett (1982), is effective and has changed little over time (Lubell and Brand, 2018). Repeated cycles of self-pollination using this technique will lead to increased homozygosity and possible inbreeding depression (Kurtz *et al*., 2020).

Floral phenology is an important component in hemp breeding programs because substantial variation can affect the uniformity of cultivar populations (Stack *et al*., 2021). Phenological descriptors for hemp are often unclear because of latitudinal variance in light conditions. Cultivars are considered to be photoperiod-sensitive when time-to-flower depends on the night length threshold, which can range from 8 to 12 hours (Hall *et al*., 2014; Moher *et al*., 2021).

These cultivars can maintain vegetative growth indefinitely if a night length of less than the critical threshold is maintained. For photoperiod insensitive cultivars (day neutral), flowering is dependent on other factors, like plant maturity. Day neutrality is advantageous at high latitudes, where the growing season is short, and at low latitudes, where daylength is insufficient to synchronize flowering. At mid-latitude growing regions, populations derived from higher latitudes begin within a few weeks after germination, while those from equatorial latitudes remain vegetative for many months. Historic latitudinal adaptation and recent admixture have led to complex epistatic interactions between the genetic factors controlling flowering in hemp. Since flowering time is of such importance for hemp yield and cultivar adaptation, there is considerable interest in identifying genes responsible for variation of the trait (Petit *et al*., 2020).

Hemp is overall more genetically diverse and heterozygous than high-THC varieties of *C. sativa* (Sawler *et al*., 2015). Substantial admixture coupled with few founders has narrowed the genetic base of drug-type germplasm, which is relatively distinct from natural hemp populations. Cannabinoid hemp cultivars bred for CBD production were derived from crossing fiber hemp with high-THC lines to take advantage of the historical selection for increased total cannabinoid content (van Bakel *et al*., 2011; Grassa *et al*., 2021). There has been relatively little analysis of the genetic diversity and heterozygosity of hemp based on founder breeding pedigrees or market class. When considering the development of hybrid cultivars of hemp, it will be important to understand whether there are heterotic groups and if there are correlations between hybrid progeny performance and genetic relatedness of the parents, as there are in maize (Marsan *et al*., 1998).

At present, there is no ideotype for cannabinoid hemp because optimized agronomic management practices and efficient harvesting equipment have yet to be established. Hemp cultivation in the US mainly uses transplants of greenhouse-grown seedlings or rooted cuttings frequently grown in plasticulture with wide row (2-3 m) and in-row (0.8-1.2 m) spacing. As the market evolves, it is likely that cannabinoid cultivation will move towards direct-seeded, high-density plantings. However, improvements in the efficiency of feminized seed production would be needed to achieve this. Nevertheless, emerging markets will continue to support small farmers and niche products, such as growing cultivars with diverse cannabinoid and terpene profiles (Andre *et al*., 2016).

While there is no consensus on how best to integrate aspects of plant architecture as selection criteria in contemporary hemp breeding programs, it has been somewhat altered by domestication (Clark and Merlin, 2016). Today, there are three major hemp market classes (Fike *et al*., 2020) that are defined by their primary end-product: grain, fiber, and cannabinoids. In general, grain hemp cultivars have been selected for early maturity and large seed size, fiber hemp for unbranched stalks and long internodes, and cannabinoid hemp for maximum flower yield, which are profusely branched and have shorter internodes. Early selection of cannabinoid hemp before terminal flowering is a challenging because the primary harvestable end-product, the inflorescence, does not form until the critical photoperiod is exceeded. If intensive early selections could be made before floral initiation, those individuals could be vegetatively propagated, then crossed to improve genetic gain, rather than making selections after pollination has occurred. The identification of a suite of traits that are evident early in plant growth and development, which are associated with desired end of season morphology, could provide informative phenotypes for indirect selection. At present, there are few experimental reports that have focused on the heritability of canopy architecture traits that might serve as useful tools for hemp breeding and selection.

The main objectives of this study were to (1) assess the genetic diversity of hemp, (2) evaluate common parent families segregating for economically-important traits in the field, (3) develop novel selection criteria for plant architecture, (4) determine important predictors of final dry floral biomass yield, and (5) establish an ideotype for field grown cannabinoid hemp.

## Materials and Methods

### DNA isolation, genotyping, and diversity analysis

Leaf tissue from 190 hemp genotypes representing grain, dual-purpose, fiber, and cannabinoid cultivars, as well as U.S. feral accessions, were obtained from Cornell Hemp seed and clonal inventories or from breeders or growers (Table S1). Young leaves and shoot-tips were freeze dried, then ground to a fine powder with a Geno/Grinder^®^ (SPEX SamplePrep, Metuchen, NJ), and genomic DNA was extracted using a DNeasy^®^ Plant Mini kit (QIAGEN Inc., Valencia, CA), following the manufacturers protocol. DNA quality was checked by agarose gel electrophoresis and quantified with a Qubit^®^ fluorometer (Thermo Fisher Scientific, Waltham, MA). Library construction and sequencing was based on the 96-plex genotyping-by-sequencing (GBS) protocol (Elshire *et al*., 2011), with ApeKI serving as the restriction enzyme. Targeting 2.5 million reads per sample, 2×150bp libraries were sequenced on the NovaSeq 6000 (Illumina Inc., San Diego, CA) platform at the University of Wisconsin Biotechnology Center DNA Sequencing Core Facility (Madison, WI).

Variant discovery was performed using the Tassel GBS Discovery Pipeline version 2 (Bradbury *et al*., 2007; Glaubitz *et al*., 2014). Barcoded reads were aligned to the CBDRx version 2 genome assembly GCF_900626175.2 (Grassa *et al*., 2021) with BWA *mem* (Li and Durbin, 2009). The mean number of barcoded reads per sample was 2.26 million with a mean alignment of 91%. The raw vcf, containing 275,828 single nucleotide polymorphisms (SNPs), was filtered with VCFtools version 0.1.15 (-minDP 7, -minQS 30, -max-missing 0.15, -maf 0.01) (Danecek *et al*., 2011), resulting in 55,452 SNPs covering all 10 chromosomes, 20 unplaced scaffolds, and the mitochondrial genome. Mean sample heterozygosity was 17% (range: 6% to 34%) and mean site missingness was 16% (range: 7% to 27%).

A distance matrix, calculated as 1 – pIBS (probability of identity-by-state), was converted to Newick format to generate an unrooted neighbor-joining tree in Archaeopteryx (Han and Zmasek, 2009). To infer the number of clusters for discriminant analysis of principal components (DAPC), variants were converted to genind format in vcfR (Knaus and Grünwald, 2017), which served as input to *find.clusters* (max.n.clust = 19, n.pca = 50, n.start = 10, n.iter = 100, method = ‘kmeans’, stat = ‘BIC’) and *dapc* in adegenet 2.0 (Jombart *et al*., 2010). Pairwise fixation index (F_ST_) and allelic richness (rarified allele counts) were estimated for each cluster using *genet.dist* (method = ‘WC84’) and *allelic.richness*, respectively, in hierfstat (Goudet and Jombart, 2021).

### Population development

Genetically diverse female hemp plants were crossed with the high-concentration cannabidiol female hemp cultivar, ‘TJ’s CBD’ (Stem Holdings Agri, Eugene, OR), to generate 17 common families and another six families were produced using two inbred S_1_ selections of ‘TJ’s CBD’ (Table 1). To ensure all progeny were female, silver thiosulfate (STS) was used to induce staminate flowers on the pollen parent (Ram and Sett, 1982). Three weekly foliar spray applications of 8 mM STS was sufficient to initiate productive staminate flowers. Crosses were conducted in separate greenhouses to ensure there was no undesired cross-pollination. Plants were dried, threshed, seed was cleaned of debris, and stored in a secure, locked freezer (−4°C) to ensure seed longevity.

**Table 1.** Cornell Hemp accession numbers, family pedigree, and inferred group of the seed parent.

| Accession | Seed parent | Pollen parent | N | Inferred group |
| --- | --- | --- | --- | --- |
| GVA-H-19-1153 | A2R4 #301 | TJ's CBD | 15 | Feral/Fiber |
| GVA-H-19-1154 | AC/DC × Otto II #4 | TJ's CBD | 15 | BaOx/Otto II |
| GVA-H-19-1157 | Candida #1 | TJ's CBD | 15 | Cherry |
| GVA-H-19-1160 | Cherry Wine #4 | TJ's CBD | 15 | Cherry |
| GVA-H-19-1161 | Cornell-OP #1 | TJ's CBD | 15 | West Coast |
| GVA-H-19-1162 | Cornell-OP #2 | TJ's CBD | 15 | T1/R4 |
| GVA-H-19-1164 | Double Cherries #2 | TJ's CBD | 15 | Cherry |
| GVA-H-19-1166 | FL 58 | TJ's CBD | 15 | Cherry |
| GVA-H-19-1169 | Otto II #3 | TJ's CBD | 15 | BaOx/Otto II |
| GVA-H-19-1170 | R4 #6 | TJ's CBD | 11 | T1/R4 |
| GVA-H-19-1171 | The Housewife #1 | TJ's CBD | 15 | West Coast |
| GVA-H-19-1174 | Otto II #3 × NEBf | TJ's CBD | 15 | BaOx/Otto II |
| GVA-H-19-1177 | R4 × Cherry Wine #8 | TJ's CBD | 15 | T1/R4 |
| GVA-H-19-1178 | R4 × Cherry Wine #10 | TJ's CBD | 15 | T1/R4 |
| GVA-H-19-1197 | NY Feral #1 | TJ's CBD | 15 | T1/R4 |
| GVA-H-19-1205 | TJ's CBD S<sub>1</sub> #4 | TJ's CBD S<sub>1</sub> #4 | 15 | West Coast |
| GVA-H-19-1206 | TJ's CBD S<sub>1</sub> #5 | TJ's CBD S<sub>1</sub> #5 | 13 | West Coast |
| GVA-H-20-1001 | Candida #1 | TJ's CBD S<sub>1</sub> #5 | 13 | Cherry |
| GVA-H-20-1028 | (PR × FD) × NEBf #1 | TJ's CBD | 15 | Feral/Fiber |
| GVA-H-20-1063 | Lifter × TJ's CBD #89 | TJ's CBD | 15 | West Coast |
| GVA-H-20-1072 | A2R4 #21 | TJ's CBD S<sub>1</sub> #4 | 13 | Feral/Fiber |
| GVA-H-20-1087 | R4 #6 × TJ's CBD #11 | TJ's CBD S<sub>1</sub> #4 | 15 | T1/R4 |
| GVA-H-20-1117 | R4 #6 × TJ's CBD #15 | TJ's CBD S<sub>1</sub> #4 | 15 | T1/R4 |

### Experimental design

For each family, seeds were sown into deep 50-cell Sureroots trays (T.O. Plastics, Clearwaterm MN) with potting mix (LM111, Lambert, Rivière-Ouelle, QC, Canada) in the greenhouse with supplemental lighting at 16:8 light:dark regimen, three weeks before planting in the field at Cornell AgriTech (Geneva, NY). The common parent, ‘TJ’s CBD’, was planted from cuttings, but grown in the same greenhouse conditions as the seedlings. Cuttings were rooted using Clonex^®^ Rooting Gel (Hydrodynamics Intl., Lansing, MI). At the time of planting (June 16, 2020), 15 progeny individuals were randomly selected from each family, and planted together in single plots at 1.2 m spacing within row and 1.8 m spacing between rows (Fig. S2). Granular fertilizer (19-19-19, N-P-K) was incorporated at 95 kg ha^-1^ before raised beds with plastic mulch were built. Drip irrigation was installed under plastic mulch. Landscape fabric was installed in aisles to suppress weed pressure. Soil moisture sensors (HOBOnet 10HS, Onset Computer Corp, Bourne, MA) were randomly installed across the field to aid in timing of irrigation. The field was fertigated twice through a Dosatron (Dosatron Intl., Inc., Clearwater, FL) four and six weeks after planting, using Jack’s 12-4-16 Hydro FeED RO (J.R. Peters Inc., Allentown, PA).

### Floral phenology

Flowering date was recorded when pre-terminal (axial flowers with shortening internodes) and terminal pistils (clusters of flowers at shoot termini) were observed. Weekly observations were recorded until all plants were terminally flowering, and expressed as days to flower after planting in the field.

### Stem growth and architecture measurements

Plant height, measured as the length of the primary stem, was assessed weekly for 10 weeks, beginning the week after planting in the field. Growth rates were calculated using height measurements over time. Plant form was derived from plant height, maximum canopy diameter and height at that diameter at 63 DAP (Table S2; Fig. S3). From these measurements, upper kite hypotenuse to lower kite hypotenuse ratio was calculated, as well as kite perimeter and kite circularity. Trunk length (ground to first branch) was measured at harvest, and was subtracted from final height to scale kite area. Internode length was derived from counting the number of branching pairs along 50 cm of the primary stem in the middle of the canopy. Kite branch angle was calculated from the lower kite triangle, using the difference of maximum canopy diameter height and trunk length to scale the hypotenuse.

### Foliar and physiology measurements

Chlorophyll concentration was measured using an Apogee MC-100 meter (Apogee Instruments Inc., Logan, UT) at 50 DAP, as an average of three unique measurements on fully-expanded leaves in the middle canopy, not less than 15 cm from the shoot apex. Leaves at the same position were sampled from the outer canopy at 86 DAP to measure leaflet number, leaf area, leaf length, leaf perimeter, petiole area, petiole length, and petiole perimeter. Intact leaves with petioles were measured in the Fiji distribution of ImageJ (Schindelin *et al*., 2012) by converting each leaf scan image to 8-bit binary then manually separating leaf and petiole. For all leaf area measurements, leaflets were attached to the rachis. Leaf width was not measured because of the non-uniform placement of leaflets in leaf scans. Alternatively, the maximum width and length of the middle leaflet was measured, and were used to derive middle leaflet area, calculated as a pointed oval. Entire leaf area (sum of leaf and petiole) was validated (R^2^=0.9993) by comparing manual measurements from ImageJ with automated measurements using *run.ij* (low.size = 0.5, low.circ = 0) in LeafArea (Katabuchi, 2015). Each leaf and respective petiole were then dried and weighed to calculate specific leaf area and specific petiole area. Using the same leaf scans, green leaf index, (2G−R−B) / (2G+R+B), was calculated by splitting and masking RGB channels in imager (Barthelme, 2017) (Fig. S4). Hemp powdery mildew (*Golovinomyces spadiceus*) severity (PM) was visually rated for each plant on a continuous scale of 0-100% canopy leaf area diseased, measured at 71, 86, and 97 DAP.

### Biomass measurements

Individuals were harvested when the inflorescence was fully-mature, typically five weeks after initiation of terminal flowering. Total wet biomass yield was measured for all individuals in the trial. For each family, a representative individual (34 individuals in total) was harvested and dried to calculate dry biomass yield, then floral material was hand stripped and weighed to determine floral biomass yield. To account for differences in growth rates attributable to flowering time variation in segregating families, representative early- and late-flowering individuals were harvested for dry biomass measurements. The wet biomass of both early and late flowering samples was strongly correlated with dry biomass (r = 0.98) and dry stripped biomass (r = 0.96). The strong linear relationship of wet to dry biomass made accurate prediction of dry stripped biomass achievable. Utilizing the wet (WBM), dry (DBM), and dry stripped biomass (DSBM) of the sampled individuals, predictions were obtained with the following simple linear models: DBM = −0.13322 + 0.31174 × WBM + ε, where ε ∼ 𝒩(0, 0.1446^2^); DSBM = 0.113884 + 0.156749 × WBM +ε, where ε ∼ 𝒩(0, 0.1031^2^). Dry floral biomass per unit area (kg m^-2^) was calculated by dividing dry floral biomass by the square of the maximum canopy diameter.

### Phenotypic characterization based on RGB and multispectral UAS images

A Matrice 100 series drone equipped with a Zenmuse 3 RGB camera (4K 4096 × 2160 px) (DJI, Shenzhen, China) and a MicaSense RedEdge five-band multispectral sensor (1280 × 960 px) (MicaSense Inc., Seattle, WA) was flown 10 times during the growing season using the DroneDeploy App version 2.90.0 (DroneDeploy, Sydney, Australia). Flights were completed at 10 day intervals from 15 DAP to 93 DAP, with an altitude of 20 m and 80% front and side overlap. Ground sampling distances for the Zenmuse 3 and RedEdge were 0.86 cm/px and 1.39 cm/px, respectively. A data processing pipeline was developed to analyze collected aerial images for the extraction of morphological and vegetation index traits (Fig. S5). Collected color and multispectral images were processed using Metashape Pro version 1.6.0 (Agisoft LLC, Russia) to reconstruct color and multispectral orthoimages and colorized 3-D point clouds. Ground control points were manually surveyed using a real time kinematic Trimble R8s GPS (Trimble Inc., Sunnyvale, CA), and used to georectify the reconstructed data in the universal transverse mercator coordinate system for successive analyses.

Plant geo-locations were calculated using color orthoimages. A color orthoimage was converted to an excessive green index (Woebbecke *et al*., 1995) map then binarized using the Otsu method (Otsu, 1979). Connected component labeling was used to segment individual plants and calculate their center locations. Based on plant centers, bounding boxes of 1.83 m (across row) and 1.22 m (within row) were generated for the localization and segmentation of plants in point clouds and multispectral orthoimages. A significant shift of plant centers was observed between 23 and 34 DAP, so the plant geo-locations and bounding boxes were derived from the color orthoimages on the two days, respectively. The locations and bounding boxes calculated on 23 DAP were used to analyze the data collected on 23 DAP, and those calculated on 34 DAP were used for the rest of data.

In the colorized point clouds, the point cloud of each plant was cropped using the calculated bounding boxes. Random sample consensus (Fischler and Bolles, 1981) was used to identify the ground plane in the plant point cloud (red points in Fig. S5) and separate canopy points (green points in Fig. S5) for the extraction of canopy morphological traits: height, projected area, and volume. In the multispectral orthoimages, a circular region with a radius of 0.28 m was defined at each plant center, and seven vegetation indices were calculated using pixels within the region for a corresponding plant. The seven vegetation indices include normalized difference vegetation index (NDVI) (Rouse *et al*., 1973), enhanced vegetation index (EVI) (Huete *et al*., 2002), green chlorophyll index (GCI) (Gitelson *et al*., 2003), green normalized difference vegetation index (GNDVI) (Gitelson *et al*., 1998), modified non-linear index (MNLI) (Yang *et al*., 2008), modified soil adjusted vegetation index 2 (MSAVI2) (Qi *et al*., 1994), and optimized soil adjusted vegetation index (OSAVI) (Rondeau *et al*., 1996). Equations for physiological indices are in Table S2.

### Cannabinoid analysis

The primary terminal inflorescence (10 cm) was sampled from each individual in the week preceding harvest for cannabinoid analysis, and dried in a climate-controlled room with a maximum temperature of 30°C and average relative humidity of 35%. Once dried, samples were milled to a fine powder for cannabinoid analysis via HPLC, further described in Stack *et al*. (2021). To control for potential variation in decarboxylation of acid-form cannabinoids, statistical analyses were based on total potential cannabinoid percentages by mathematically combining the concentrations of the acid and neutral form as described in Stack *et al*. (2021).

### Statistical analysis

All plotting and statistical analyses were conducted in the open-source statistical computing platform R version 3.4.2 (R Core Team, 2017). Following ANOVA, mean separation was conducted using Tukey’s HSD in agricolae (Mendiburu, 2017). Wilcoxon Rank Sum test was used for paired comparisons with *wilcox.test* (conf = 0.95). Tests for pairwise associations (p < 0.001) were conducted using Pearson’s correlation coefficient (r) with *cor.test*. Model II linear regression was conducted using the ranged major axis (RMA) method in lmodel2 (Legendre, 2014). Maximum growth rates were modelled with *all_splines* (spar = 0.35, optgrid = 50) from the log-linear part of the growth curve in growthrates (Petzoldt, 2007). Absolute area under the disease pressure curve (AUDPC) was calculated from three PM ratings, using the *audpc* function in agricolae. Archetypal analysis of plant architecture was performed with *stepArchetypes* (k = 1:10, nrep = 5) in archetypes (Eugster and Leish, 2009), using the two ratios as input variables: maximum canopy diameter to plot height and maximum canopy diameter height to plot height. Heritability was calculated as the ratio of additive genetic variance to total phenotypic variance. For half-sib families, additive genetic variance was estimated as four times the family variance (σ^2^_A_ = 4σ^2^_F_) and phenotypic variance as the sum of family and residual variance. Variance components were estimated using *lmer* in lme4 (Bates *et al*., 2015). Variable selection via stepwise regression was performed with *stepAIC* (direction = ‘both’). Relative importance metrics for multiple linear regression was performed to order predictors and decompose R^2^ in relaimpo (Grömping, 2006), using 1000 bootstrap replicates (Bonferonni CI = 95%). LMG indices (Lindeman *et al*., 1980) were used to partition additive properties of R^2^, calculated as the sum of their individual importance, irrespective of the correlation among predictor variables.

## Results and Discussion

The relationships between architecture, foliar, physiological, and pathology traits, and HTP measurements were assessed to provide a clearer picture of hemp growth and development, and to understand their contribution to the traits of primary economic importance: floral biomass and cannabinoid yield. These were assessed in families with a common parent to reveal the degree of segregation based on parent heterozygosity and allelic diversity.

### Genetic diversity of hemp

Parent selection for this study was predicated on the genetic relatedness of a broad sample of available hemp germplasm using high-density SNP markers (Table S1), of which seed parents from all cannabinoid hemp clades are fully representative (Fig. 1A). Of the seven primary clusters identified, genotypes grouped by market class (grain, fiber, cannabinoid), geographical origin, and/or a common founder (Fig. 1B; Fig. 1C). For instance, grain, Italian fiber and U.S. feral, and Chinese genotypes were represented by distinct clusters. Cannabinoid hemp genotypes clustered by common founder populations and were the most admixed; however, cannabinoid genotypes in group 1 (T1/R4) were particularly distinct, with no clear evidence of admixture (Fig. 1D). There was clear population differentiation, with pairwise F_ST_ ranging from 0.08 (grain/dual and fiber/feral) to 0.30 (grain/dual and T1/R4) (Fig. 1E), reflecting the spatial separation of DAPC clusters in Fig. 1B. Increased allelic richness was observed at the end of chromosome 4 for the fiber/feral cluster (Fig. S1), and all four cannabinoid clusters had lower allelic richness along chromosome 7, probably due to repeated selection on cannabinoid synthase cassettes residing therein (Grassa *et al*., 2021).

**Fig. 1.**
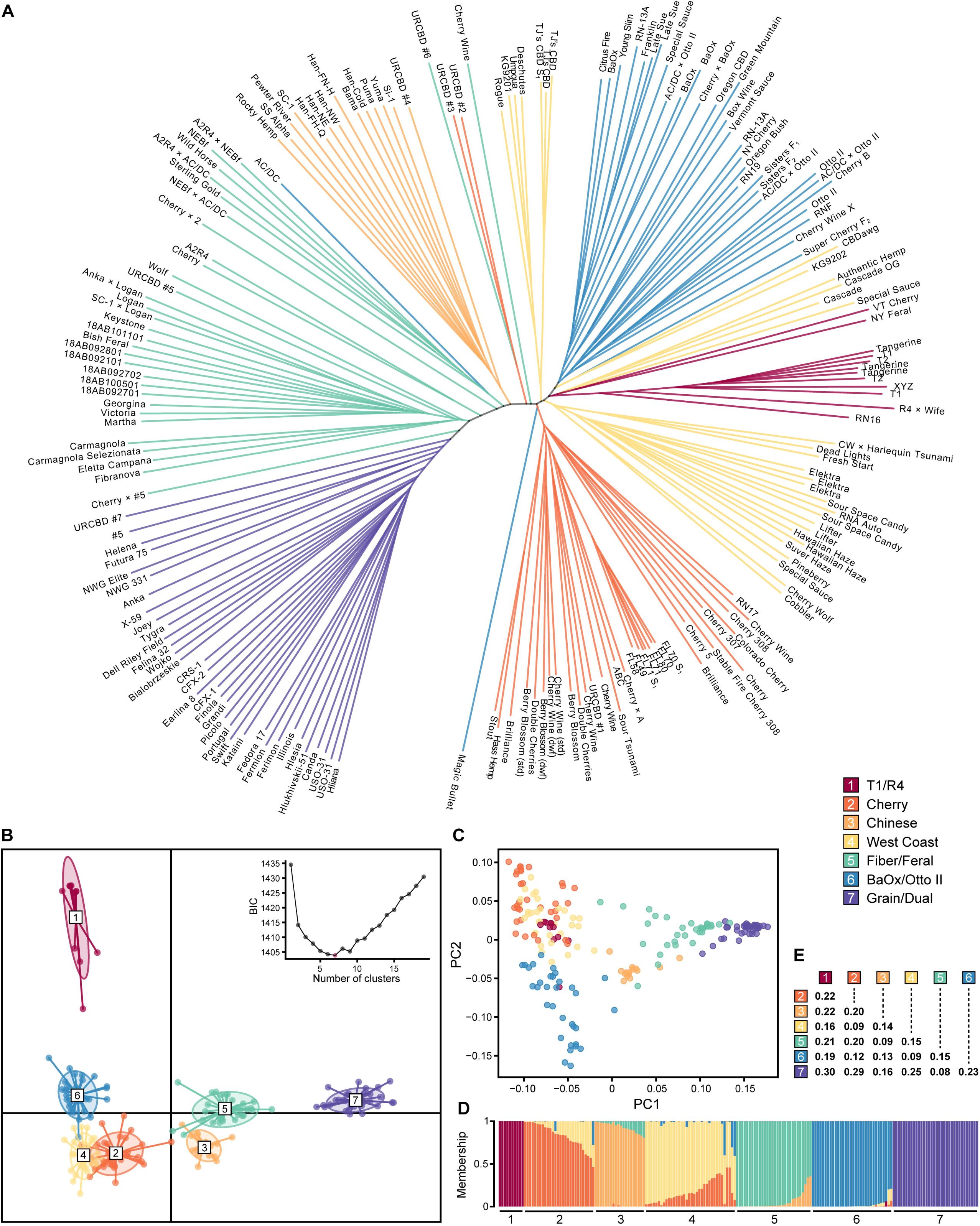
Genetic diversity of hemp. Using 190 hemp cultivars, crosses, and feral accessions, panels depict **(A)** unrooted neighbor-joining tree, **(B)** discriminant analysis of principal components (DAPC), **(C)** principal component analysis, **(D)** membership probability ordered by DAPC cluster and within-cluster membership, and **(E)** pairwise F_ST_. DAPC clusters 1-7 inferred via Bayesian Information Criterion (BIC) (k = 1:19) are labelled/colored according to the legend.

Hybrid development in hemp will depend on genome-wide markers to infer kinship and heterotic group membership. Prediction of combining ability is the hallmark of hybrid breeding, however, a deficiency of published studies on hemp hybrids and the lack of a unified genotyping platform has delayed its progress in the research community. Nevertheless, the clusters identified here serve as a good starting point for hemp breeders to select representative inbred parent lines for early analysis of combining ability and stable hybrid deployment.

### Floral phenology is quantitative with major effect genes

There was substantial variation in flowering time both within and among families (Fig. 2A). Seven families segregated for early, mid, and late terminal flowering day, of which earlier flowering individuals were far less variable compared to those flowering later. The mean number of days to pre-terminal and terminal flowering was 59.3 DAP (range: 28 to 85 DAP) and 68.9 DAP (range: 35 to 92 DAP), respectively. The common parent clone, ‘TJ’s CBD’, initiated flowers at 63 DAP and terminal flowering at 70 DAP, and did not vary across replicate plots. While floral phenology in *C. sativa* is quantitative (Salentijn *et al*., 2019), there may be few genes conferring the early flowering trait observed in this population. Based on the segregation of the S_2_ families, the considerable within-family variation in days to flower was due to the common parent being heterozygous for at least one gene of major effect in the flowering time pathway. For instance, S_2_ family 19-1206 initiated flowering at 42 DAP, whereas, in S_2_ family 19-1205, eight progeny initiated at 42 DAP and seven after 63 DAP. Family 20-1001, which had the same S_1_ pollen parent as S_2_ family 19-1206, flowered the earliest at 35 DAP. In addition, the seed parent of 20-1001 ‘Candida’ was also crossed with the original heterozygous common parent (family 19-1177), with five terminally flowering ∼35 DAP and 10 ∼85 DAP.

**Fig. 2.**
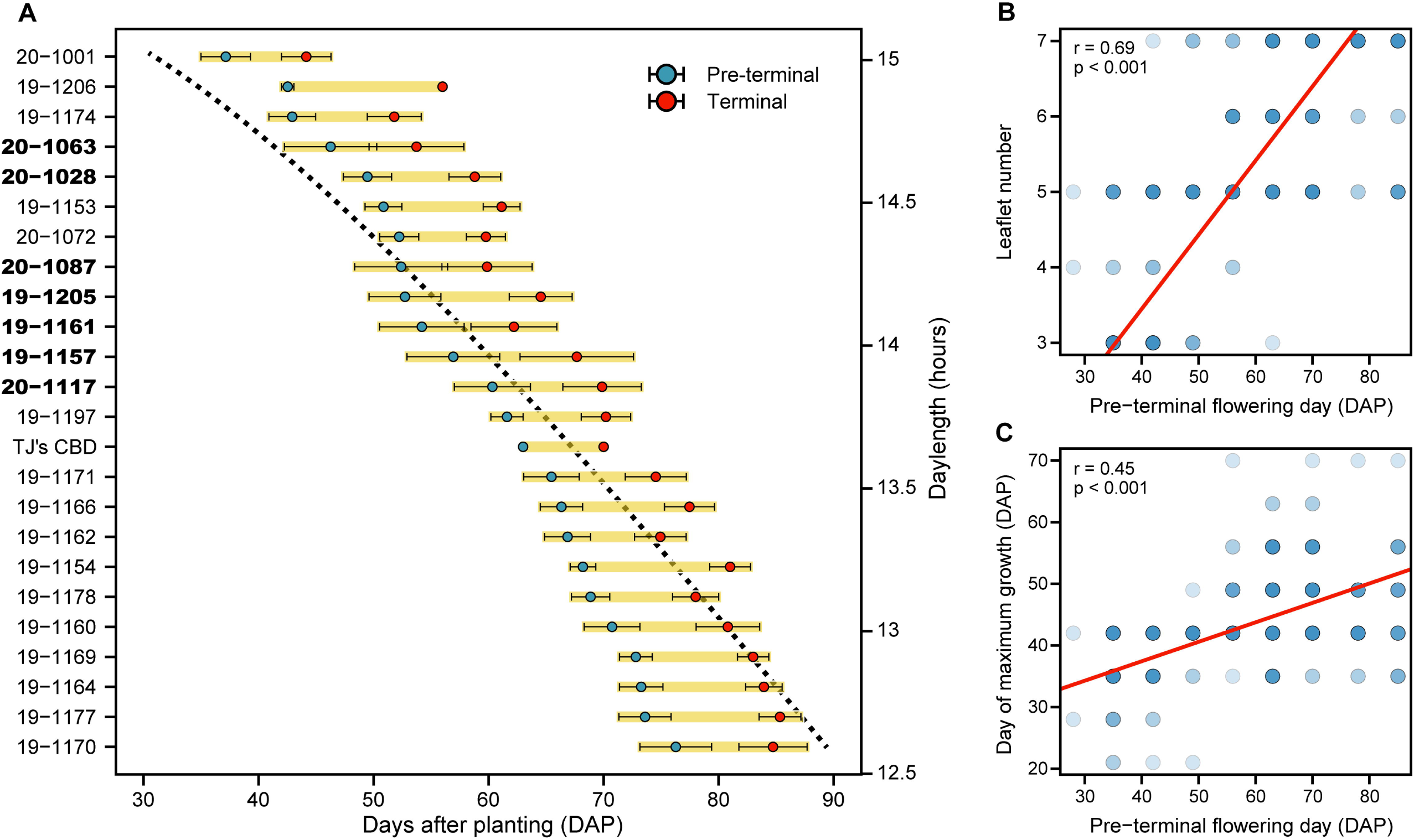
Floral phenology. Ordered by mean pre-terminal flowering day, **(A)** family means (± standard error) for pre-terminal (blue points) and terminal (red points) flowering. The black dotted line represents daylength over the same time period on the alternate y-axis. Bold family names represent those families segregating for flowering date. Regression of **(B)** leaflet number and **(C)** day of maximum stem growth with pre-terminal flowering day, each represented in days after planting (DAP).

For those families not clearly segregating for flowering date, the presumably dominant seed parent allele(s) masked that of the common parent in the hybrid. If this was a simple recessive trait, S_1_ progeny would segregate 3 late : 1 early, and depending on the status of the S_1_ parent, S_2_ lines would be either all-early, all-late, or segregating. However, 3:1 ratios were not clearly observed, but were either ∼1:1, ∼2:1, all-early, or all-late. If 1:1 ratios could be explained as a testcross, then homozygous recessive seed parents would express the early flowering trait, yet only ‘Candida’ and ‘TJ’s CBD S_1_ #5’ flowered early. It is probable that more than one gene is controlling the early flowering trait observed in this population, and may be evidence of epistasis. Since the seed parents were genetically diverse, there are likely to be multiple segregation models for this trait. Although some ratios could be explained by a single factor and others, two or more, larger populations would undoubtedly be required to test these hypotheses. Inbred lines can be developed that will be homozygous for those major gene alleles affecting flowering time, but it will be critical to understand how different alleles interact in the heterozygous state in F_1_ hybrid cultivars to determine latitudinal adaptation.

### Foliar allometry

Leaf morphology traits are routinely used to designate plants as *C. sativa* (narrow leaflets), *C. indica* (broad leaflets), *C. ruderalis* (three leaflets to a leaf), or hybrids and/or “percentages” of each (Clarke and Merlin, 2013), albeit weakly justified (Vergara *et al*., 2021). Here, fully-expanded leaves were collected from each individual at the same approximate location, irrespective of terminal flowering day. All three general leaf types could be observed on individual genotypes. These observations can be attributed to ontogenetic heterophylly, notwithstanding confounding factors like variation in floral phenology and canopy light penetration. There was a strong positive correlation of terminal flowering day and leaflet number (r = 0.67) (Fig. 2B), of which three primary groups of individuals had a common terminal flowering day, and mean leaflet number increased with these groups: <50 DAP (3.9±0.14), 50-70 DAP (5.3±0.09), and >70 DAP (6.3±0.09). Since early flowering individuals formed inflorescences at shoot apices early on, it is possible that differential levels of a growth hormone, such as ethylene, may have had an effect on leaf development and morphology, compared to those flowering later.

The mean leaf and petiole area for families with seed parents inferred to be in T1/R4 diversity group 1 (Table 1) were 37% to 54% and 34% to 82% greater than mean values of all other family groups, respectively. With a population mean of 7.8 cm^2^ leaflet^-1^, families 19-1162, 19-1170, and 19-1197 (all in: T1/R4 diversity group 1) had the greatest mean leaflet area (>10.2 cm^2^), whereas families with the lowest mean leaflet area were 20-1072 and 19-1153 (<4.9 cm^2^), both of which shared the common seed parent ‘A2R4’ (all in: Fiber/Feral diversity group 5). In general, families with a Fiber/Feral seed parent background had smaller leaves and petioles, and those with a T1/R4 background were most always larger.

Larger plants were apt to have a lower leafing intensity (r = −0.82), which is defined as the ratio of the number of leaves to stem volume. It is thought that smaller leaves of land plants are found on species that produce more of them, such that variation in leaf size can be predicted in terms of a leaf mass/number trade-off. Kleiman and Aarssen (2007) found that in the new growth of deciduous trees, the slope of log (leafing intensity) and log (leaf mass), does not significantly deviate from −1. They posit that selection may favor high leaf intensity, with small leaf mass resulting not as direct adaptation, but simply as a trade-off. Here, the slope of the regression of log-transformed leafing intensity and leaf mass (R^2^ = 0.66) was −0.71 (Fig. 3A), indicating that for every 1% increase in leafing intensity, there was a 0.71% decrease in individual leaf mass. Although the obvious difference here is that *C. sativa* is an herbaceous annual, calculation of total leaf number in this study included the inflorescence, which is substantial in cannabinoid cultivars. Further, cannabinoids can account for more than 20% of the dry mass of the inflorescence in mature plants, so it is very likely that the comparative decrease in individual leaf mass is an underestimate compared to that of sampled mid-season growth.

**Fig. 3.**
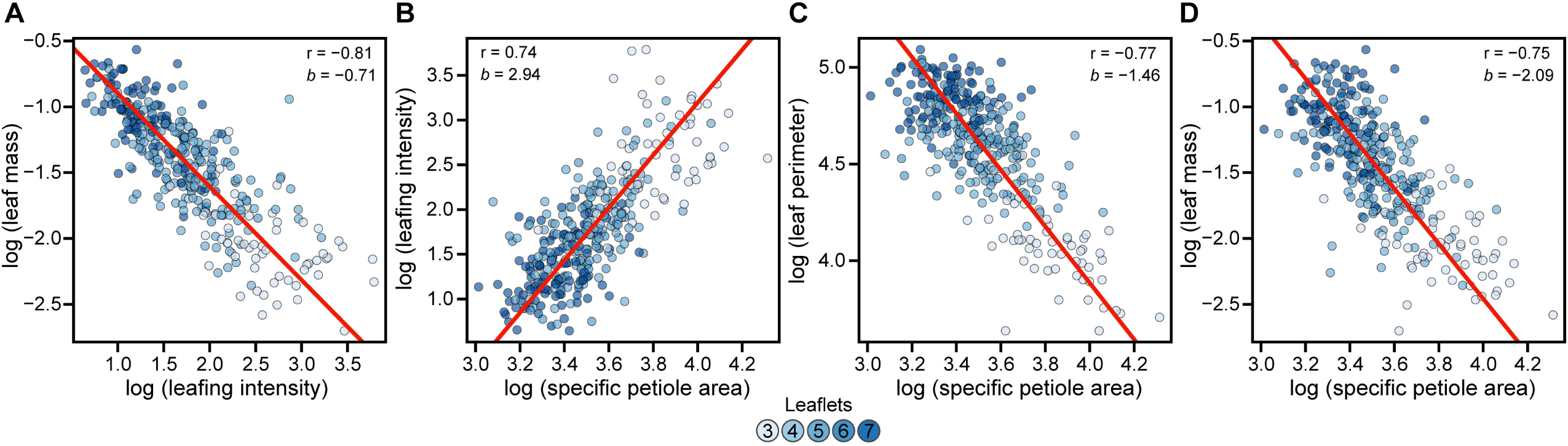
Foliar relationships. Model II linear regression of log transformed **(A)** leafing intensity and leaf mass, **(B)** specific petiole area and leafing intensity, **(C)** specific petiole area and leaf mass, and **(D)** specific petiole area and leaf perimeter. For all panels, points are colored according to the number of leaflets, as shown in the legend.

In addition, there was a strong positive relationship between leafing intensity and specific petiole area (R^2^ = 0.55, slope = 2.08), but not specific leaf area (p = 0.97) (Fig. 3B). Theoretically, specific leaf area should scale linearly with growth rate (Liu *et al*., 2021), but was not evident in this dataset (p = 0.62). Specific petiole area was inversely correlated with biomass accumulation (r = −0.51), stem volume (r = −0.49), and kite area (r = −0.48). While most leaf, stem, and physiological traits had inverse relationships with specific petiole area, petiole circularity (r = 0.23), leaf circularity (r = 0.57), and mid-canopy branch number (r = 0.44) were positively associated with the trait. An increase in specific petiole area was associated with a decrease in leaf perimeter (r = −0.77). Further, the slope of the regression of log (specific petiole area) and log (leaf perimeter) was −1.46 (R^2^ = 0.59), which suggests an allometric relationship (Fig. 3C). Similar results were found for specific petiole area and leaflet number (R^2^ = 0.52, slope = −0.9), such that plants with few leaflets per leaf had less mass per unit area and lower leaf perimeter. However, the slope of the regression of log (specific petiole area) and log (leaf dry weight) was −2.1 (R^2^ = 0.56), whereby a small increase in petiole mass per area accompanied a disproportionate increase in leaf dry mass (Fig. 3D).

### Plant growth is most influenced by floral phenology

Within-family variation in growth rate and biomass accumulation are intrinsically linked to variation in floral phenology. Overall, day of maximum stem growth was positively correlated with days to flower (r = 0.45) (Fig. 2C); however, the growth rate of all individuals did not immediately diminish following floral initiation. Families such as 19-1153, developed clusters of flowers at shoot axes and apices early in the growing season but did not begin to form compact inflorescences until weeks later. This family was also the tallest (2.05 m) and had the fastest mean growth rate (2.75 cm day^-1^) of all other families (Table S3). Not significantly different from one another, inbred S_2_ families, 19-1205 and 19-1206, had the slowest mean growth rates of 1.46 and 1.03 cm day^-1^, respectively. Furthermore, families with greater variances for final height (70 DAP) were also those segregating for early and later flowering dates.

### Kite variables are good indicators of within-family variance in canopy architecture

Kite variables produced a useful model that accurately portrays the major architectural differences in hemp. In high-density plantings of fiber and grain hemp, apical dominance is markedly stronger, and leads to a far more columnar habit that maintains dormancy of axial buds, which often results in a significant reduction of branching. The wide planting density in this study was chosen to maximize light availability so that the variation in architectural traits would be more easily discernable. The common parent to the F_1_ families in this study has a somewhat irregular, excurrent habit, which may be due to the fact that it was propagated from cuttings. Of all the individuals surveyed, not one lacked opposite decussate branching leading to sub-opposite branching at shoot apices and lateral spirals on branches.

Plant size (kite area) (Fig. 4A) and form were derived from three measurements (plant height, maximum canopy diameter, and the height at that diameter) and used to construct 2-D kite models, of which there was significant variation within and among families (Fig. 4B; Table S3). These values also displayed high heritability values, suggesting that selection will lead to genetic gains (Table 2). These kite models can be used to evaluate the segregation of plant size and form within and among populations. While there was considerable variance within some families for kite area, others were quite uniform, reflecting the low-level of heterozygosity for those seed parents. Based on archetype analysis, four primary groups (extremes) were established (Fig. 4C), each varying in kite hypotenuse ratio and/or the ratio of maximum canopy diameter to height. One form that was not observed in this study was a strongly prostrate habit (canopy diameter » height), although such an archetype would be far less economically valuable for the grower. It would be useful to understand how canopy architecture variables are altered by archetype when planted at varying densities, especially with respect to cultivar-specific suitability and environmental plasticity.

**Table 2.** Trait abbreviations, descriptions, units, as well as population means, range (min -max), coefficient of variation (CV), and heritability (h^2^). For field-collected traits with repeated measurements, the final measurement is reported. The common parent was not included in population level statistics.

| Abbreviation | Description | Units | Mean | Range | CV | $h^2$ |
| --- | --- | --- | --- | --- | --- | --- |
| *Architecture* |  |  |  |  |  |  |
| HT | Plant height | cm | 157.7 | 78 - 255 | 0.19 | 0.83 |
| BPAIR | Branching pairs in canopy | # | 7.45 | 4 - 16 | 0.26 | 0.72 |
| INL | Internode length | cm | 7.07 | 3.1 - 12.5 | 0.21 | 0.63 |
| MCD | Maximum canopy diameter | cm | 125.9 | 61 - 179 | 0.19 | 0.83 |
| MCDH | Height at max canopy diameter | cm | 85.9 | 40 - 160 | 0.24 | 0.73 |
| TRKL | Trunk length to first branch | cm | 12.2 | 1 - 50 | 0.70 | 0.60 |
| KITE | Kite area | m<sup>2</sup> | 0.94 | 0.19 - 2.14 | 0.32 | 0.82 |
| KHR | Kite hypotenuse ratio | - | 0.97 | 0.56 - 1.67 | 0.22 | 0.36 |
| KBA | Kite branch angle | ° | 41.2 | 19.9 - 67.2 | 0.16 | 0.45 |
| KC | Kite circularity | 0-1 | 0.76 | 0.57 - 0.79 | 0.03 | 0.56 |
| DIA | Basal stem diameter | cm | 4.44 | 1.6 - 6.9 | 0.23 | 0.85 |
| VOL | Primary stem volume | cm<sup>3</sup> | 888.4 | 54.9 - 1989 | 0.23 | 0.81 |
| *Foliar* |  |  |  |  |  |  |
| LFLTN | Leaflet number | # | 5.37 | 3 - 7 | 0.25 | 0.80 |
| MLFW | Middle leaflet width | cm | 1.70 | 0.89 - 2.89 | 0.21 | 0.84 |
| MLFA | Middle leaflet area | cm<sup>2</sup> | 16.4 | 7.2 - 32.3 | 0.28 | 0.79 |
| LFDW | Leaf dry weight | g | 0.25 | 0.07 - 0.57 | 0.41 | 0.84 |
| LFA | Leaf area | cm<sup>2</sup> | 42.3 | 11.5 - 97.8 | 0.41 | 0.84 |
| LFL | Leaf length | cm | 14.3 | 10.8 - 20.0 | 0.11 | 0.62 |
| LFP | Leaf perimeter | cm | 102.1 | 37.9 - 162.7 | 0.27 | 0.80 |
| SLA | Specific leaf area | cm<sup>2</sup> g<sup>-1</sup> | 167.2 | 122 - 219.3 | 0.10 | 0.59 |
| PTDW | Petiole dry weight | g | 0.03 | 0.004 - 0.08 | 0.54 | 0.80 |
| PTA | Petiole area | cm<sup>2</sup> | 0.82 | 0.2 - 2.2 | 0.43 | 0.79 |
| PTL | Petiole length | cm | 3.86 | 1.2 - 8.0 | 0.32 | 0.73 |
| PTP | Petiole perimeter | cm | 9.06 | 3.36 - 18.04 | 0.30 | 0.72 |
| SPA | Specific petiole area | cm<sup>2</sup> g<sup>-1</sup> | 34.8 | 20.4 - 74.8 | 0.24 | 0.84 |
| *Phenology* |  |  |  |  |  |  |
| PTFD | Pre-terminal flowering day | DAP | 59.3 | 28 - 85 | 0.24 | 0.85 |
| TFD | Terminal flowering day | DAP | 68.9 | 35 - 92 | 0.22 | 0.84 |
| *Physiology* |  |  |  |  |  |  |
| CCI | Chlorophyll content index | - | 51.2 | 34.8 - 78.3 | 0.15 | 0.81 |
| GLI | Green leaf index | - | 0.12 | 0.08 - 0.19 | 0.14 | 0.70 |
| GR | Mean growth rate | cm day<sup>-1</sup> | 1.90 | 0.43 - 3.4 | 0.24 | 0.80 |
| DMG | Day of maximum growth | DAP | 43.5 | 21 - 70 | 0.17 | 0.47 |
| *Pathology* |  |  |  |  |  |  |
| PM | Powdery mildew severity | % | 37.8 | 1 - 95 | 0.52 | 0.67 |
| AUDPC | PM disease progress | AUDPC | 957.8 | 18.5 - 2433 | 0.55 | 0.69 |
| *Biomass Yield* |  |  |  |  |  |  |
| WBM | Whole plant wet biomass | kg | 6.31 | 1.22 - 13.1 | 0.35 | 0.79 |
| DBM | Whole plant dry biomass | kg | 1.84 | 0.28 - 3.95 | 0.39 | 0.80 |
| DSBM | Dry stripped floral biomass | kg | 1.11 | 0.27 - 2.15 | 0.32 | 0.81 |
| *Cannabinoid Content* |  |  |  |  |  |  |
| THC | Tetrahydrocannabinol | % | 1.19 | 0.14 - 9.72 | 1.53 | 0.82 |
| CBD | Cannabidiol | % | 8.73 | 0.14 - 16.8 | 0.32 | 0.86 |
| CBC | Cannabichromene | % | 0.52 | 0.09 - 2.03 | 0.57 | 0.84 |
| CBG | Cannabigerol | % | 0.27 | 0.03 - 0.70 | 0.41 | 0.74 |
| THCV | Tetrahydrocannabivarin | % | 0.03 | 0.00 - 0.42 | 2.24 | 0.72 |
| CBDV | Cannabidivarin | % | 0.16 | 0.01 - 3.79 | 2.43 | 0.76 |
| CBL | Cannabicyclol | % | 0.04 | 0.00 - 0.12 | 0.70 | 0.70 |
| *Cannabinoid Yield* |  |  |  |  |  |  |
| CB<sub>yield</sub> | Total cannabinoids/100 × DSBM | g | 120 | 25 - 320 | 0.43 | 0.87 |
| CBD<sub>yield</sub> | Total CBD/100 × DSBM | g | 100 | 2 - 290 | 0.51 | 0.90 |

**Fig. 4.**
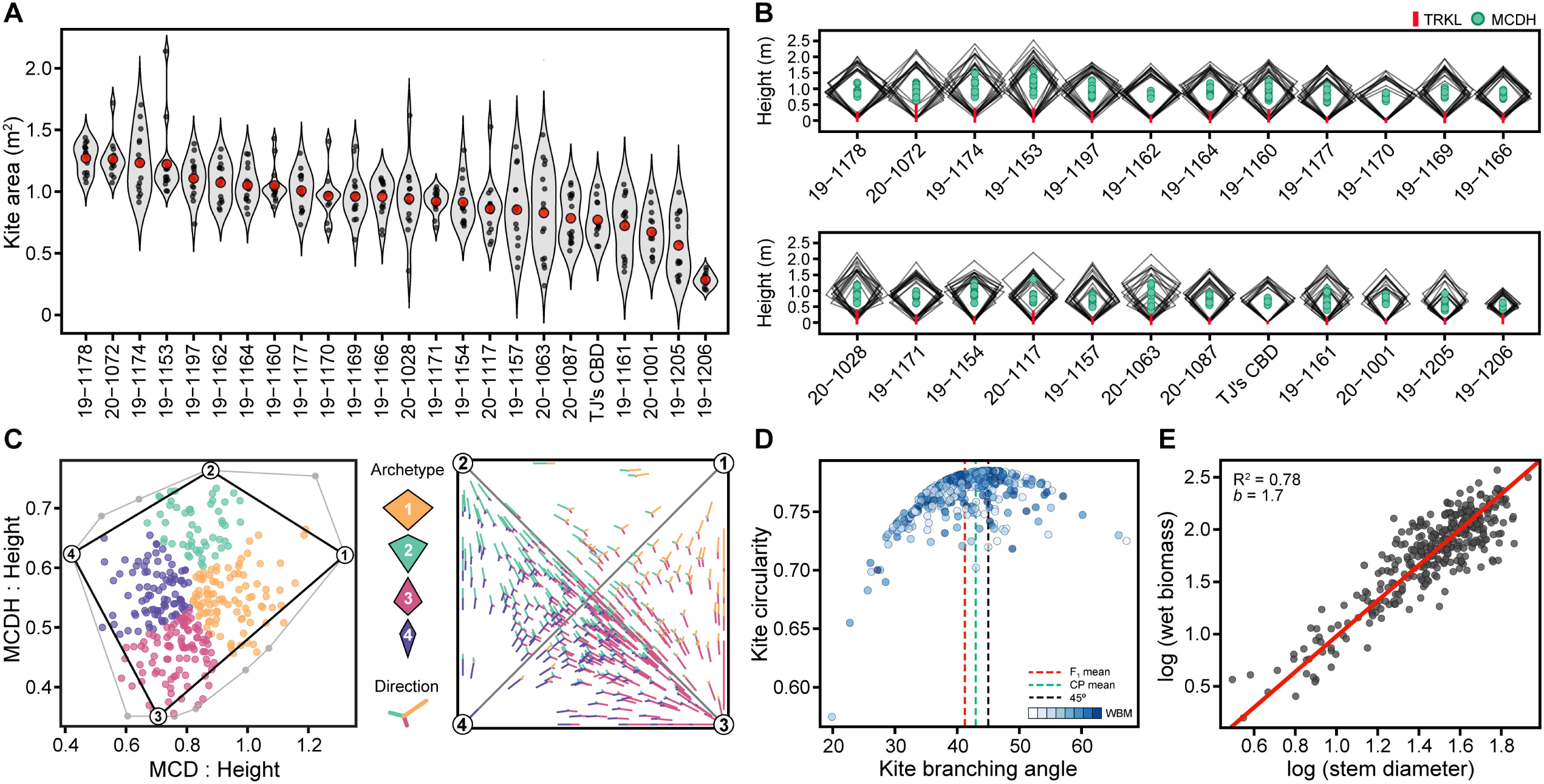
Canopy architecture. Violin plots of **(A)** kite area (m^2^) by family and **(B)** 2-D kite models in descending order. Mean canopy diameter height (MCDH) within 2-D kites are represented by filled circles and maximum trunk length (TRKL) by red lines. Axes of 2-D kites are 1:1. Archetype analysis **(C)** using the ratios of maximum canopy diameter (MCD) (x-axis) and maximum canopy diameter height (MCDH) (y-axis) to plant height, and simplex plot showing 2-D projected direction, with archetypes 1-4 depicted graphically in the legend. Scatterplot **(D)** of kite circularity and kite branch angle (KBA). Mean KBA for all F_1_ individuals, the common parent (CP), and KBA at maximum circularity (45°), are depicted in red, green, and black hashed lines, respectively. Point color opacity represents biomass accumulation, low to high. Model II linear regression of **(E)** log transformed wet biomass and basal stem diameter.

The least variable architectural trait, kite circularity, had an F_1_ mean of 0.76 (CV = 0.03) and common parent mean of 0.77. Kite circularity decreases when branch angles are either less or greater than 45°, which results in a more columnar or prostrate branching habit, respectively (Fig. 4D). While there was no significant association with kite circularity and dry floral biomass yield on a per plant basis (p = 0.4), selection of high-yielding individuals with low branch angles and low kite circularity would permit denser plantings without sacrificing axial branches to competition, and result in greater yield per acre. For instance, if dry stripped floral biomass yield is considered per unit area (kg m^-2^), both circularity (r = −0.27, p < 0.001) and branch angle (r = −0.28, p < 0.001) become redeemable selection criteria.

Since low specific kite area (m^2^ kg^-1^) is indicative of small plants with a high dry biomass proportion, and vice versa, this measurement can generally be used to infer canopy density. Theoretically, wet leaf mass fraction divided by kite volume is a more pure estimate of canopy density (kg m^-3^), and can be derived allometrically (Table S2). Canopy density was most positively correlated with the number of mid-canopy branches (r = 0.70), leaf circularity (r = 0.43), and specific petiole area (r = 0.35), and inversely with final height (r = −0.82). The family with the greatest canopy leaf density was the inbred, early-flowering S_2_ family 19-1206. Notably, the largest plants were not necessarily the most productive per unit area and dense canopies can come with significant costs, such as greater incidence of disease. Genetic selection on canopy architecture and canopy density traits described here could dramatically reduce grower inputs associated with pruning and maintenance, which has been shown to increase uniformity of cannabinoid profile in greenhouse grown, drug-type *C. sativa* (Danziger and Bernstein, 2021).

Incidents of broken branches all occurred at collars along the main stem and were caused by lateral movement from strong winds. Many broken branches continued to support foliage and inflorescences after breakage but were less vigorous. Plants with greater kite area and longer internodes were more prone to breakage, especially if branching angles were shallow. Inbred S_1_ families showed no signs of broken branches throughout the trial, due to both their small stature and shortest mean internode length (4.1 cm). However, S_2_ family 19-1206 had the greatest mean branching angle (48.7°), which suggests that plant size (kite area) plays a more important role in maintaining branch and internode lengths, which scale linearly. Besides broken branches, there were only two instances of complete lodging, each derived from the seed parent ‘A2R4’, and were significantly taller (>2 m) overall.

### Basal stem diameter is the best single predictor of biomass yield

Ordinary least square and model type II regression protocols indicate that most of the principal morphometric variables of interest, e.g., plant height, leaf size, and canopy spread, are significantly correlated (r > 0.65, p < 0.005) with basal stem diameter (Fig. S7). These findings resonate with scaling relationships reported for interspecific comparisons (Enquist and Niklas, 2002). For example, plant height, on average, scales as the 0.65-power of basal stem diameter. The numerical value of this scaling exponent is consistent with elastic self-similarity, i.e., plant height scales as the 2/3-power function of stem diameter (Niklas, 1995). In summary, most of the morphometric variables of interest are correlated with basal stem diameter (across phenotypes) and most of these variables scale with respect to one another in a manner that is consistent with scaling relationships reported for vascular plants with self-supporting stems.

Of all traits measured, basal stem diameter offers the best return on investment as a selection criteria for biomass yield. Regression of log (biomass) and log (stem diameter) resulted in a slope of 1.7 (R^2^ = 0.78), whereby incremental gains in stem diameter equate to considerable gains in biomass yield (Fig. 4E). Together with height, primary stem volume can be estimated, which is also strongly associated with biomass yield (R^2^ = 0.82, slope = 0.64), but the slope of the regression reiterates that stem diameter alone is a superior selection criterion. It would be interesting to determine the earliest point at which stem diameter measurements are predictive of biomass yield, as such information could be used in early seedling selection of breeding populations.

### Early HTP measurements are well-correlated with field phenotypes

There were dramatic differences in morphological HTP aerial measurements (canopy height, area, and volume) between flights flown before and after 56 DAP, with good correlations among measurements within but not among earlier and later flights. These differences were due to a strong wind storm between 50 DAP and 56 DAP that resulted in moderate lodging and stem breakage. Even though F_1_ families were planted in rows, a family-level analysis did not have a major effect on HTP to field phenotypic correlations of later flights (Fig. S6). Canopy height and volume obtained from orthomosaic mesh layers were well-correlated with corresponding field-collected phenotypes plot height (r = 0.83) and kite volume (r = 0.67) for early flights. Family-level correlations were even stronger for height (r = 0.95) (Fig. 5A) and volume (r = 0.80) (Fig. 5B). Biomass yield was most associated with canopy volume (35 DAP) (r = 0.56), yet this correlation was only marginally improved on a family mean basis (Fig. 5C), and for all aerial surveys beyond 50 DAP, there were only weak correlations between the two.

**Fig. 5.**
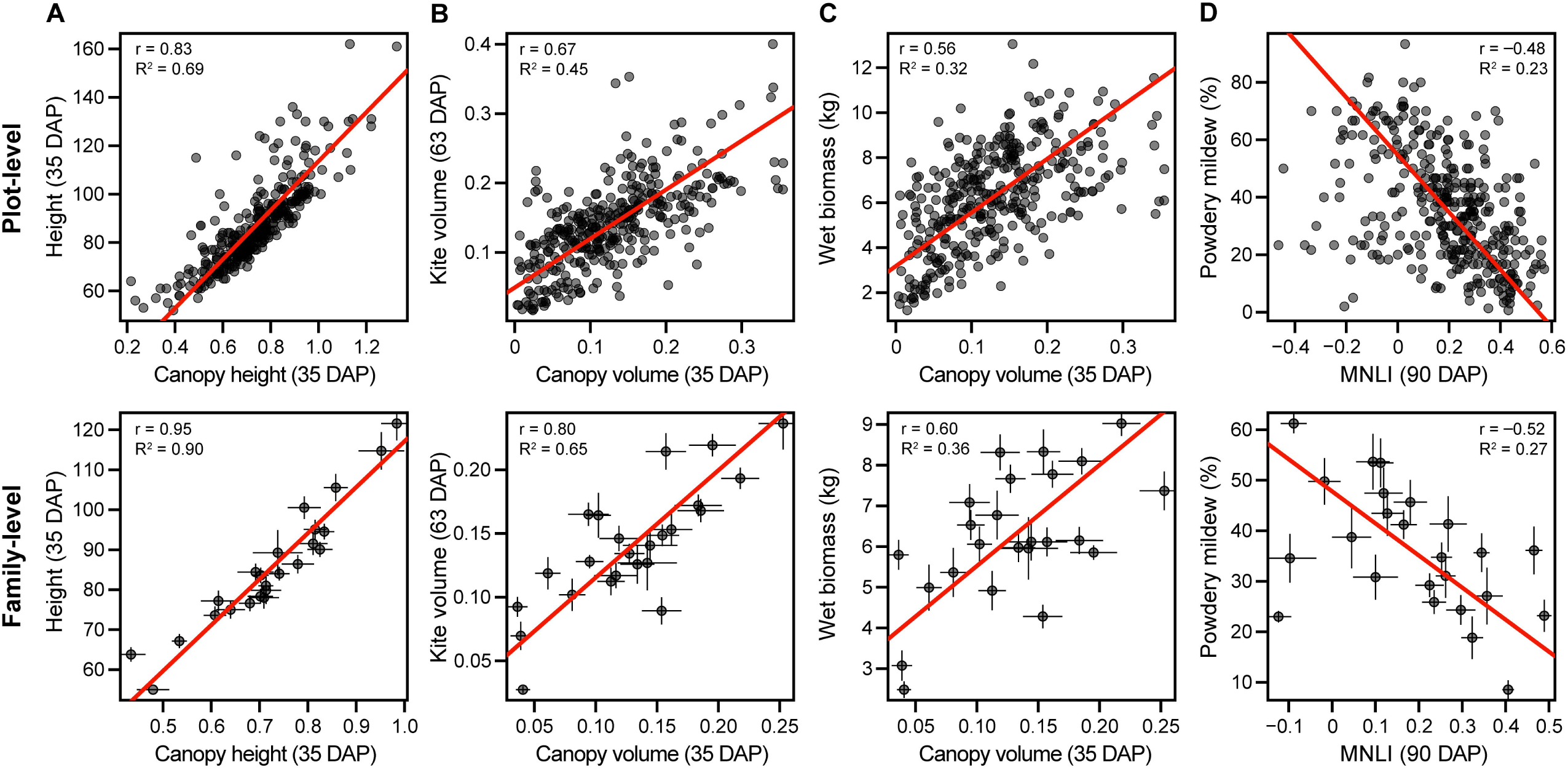
Relationships of field phenotypes and aerial indices. For panels **(A-D)**, the top figure is plot-level regression and the lower figure is family-level (mean ± standard error). Regression of **(A)** kite volume and canopy volume, **(B)** height and canopy height, **(C)** biomass yield and canopy volume, and **(D)** powdery mildew severity and modified non-linear index (MNLI).

Instances of lodging did not affect vigor or productivity but confounded the accuracy of morphological indices after 56 DAP because of alterations in the primary axis and projected area of individual plots. Physiological indices were likewise affected, but not as profoundly as the morphological indices (Fig S7). There were good phenotypic correlations with nearly all HTP measurements but EVI, which was not informative. Notably, we observed that few cannabinoids were associated with physiological indices (Fig. S8). The strongest were in the abundance of the minor cannabinoids CBL (r = −0.35) and CBDV (r = −0.17) with MNLI, MSAVI2, and OSAVI indices at 93 DAP, but those with CBDV may due to population structure, since only two families had individuals with >1% CBDV content. It may be possible to predict cannabinoid profiles and yield using multispectral or hyperspectral data, similar to what has been attempted with FT-NIR (Callado *et al*., 2018), but concerted segmentation of inflorescences would be required to develop an effective strategy to better estimate these profiles from aerial imaging. Further analyses of denser, direct-seeded plantings would both reduce incidence of lodging and offer better estimates compared to the larger plot spacing provided in this trial.

### Powdery mildew susceptibility is multigenic and affects biomass yield and quality

Biotic and abiotic factors can influence hemp yield, uniformity, and stability (Thiessen *et al*., 2020). One of the most significant diseases of hemp in the northeastern US, hemp powdery mildew (PM) is caused by the obligate biotrophic fungal pathogen *Golovinomyces spadiceus* (Szarka *et al*., 2019; Weldon *et al*., 2019), and can lead to early leaf drop and reduced inflorescence quality (Punja *et al*., 2019; Stack *et al*., 2021). Management options are available but few synthetic chemicals have been approved for field grown hemp (Punja and Scott, 2020), and genetic sources of durable resistance to PM have not been established. ‘FL 58’ (Sunrise Genetics, LLC), seed parent of family 19-1166, was previously characterized as having the lowest susceptibility to PM in multi-environment cultivar trials (Stack *et al*., 2021). In this current study, family 19-1166 displayed varying signs of PM throughout the growing season, which suggests that resistance is multigenic and/or recessive, heterozygous in the common parent background. Although there was substantial variation in PM disease progression (AUDPC) (CV = 0.55), the inherent link between floral phenology and disease incidence (i.e., early flowering plants tend to be more diseased), can make phenotypic selections for genetic resistance difficult.

Powdery mildew severity at 97 DAP ranged from 0 to 95% with a mean of 37.9% (CV = 0.52). Families with the greatest mean AUDPC were those crossed with an inbred selection of the common parent and/or derived from a selection of ‘Candida’, a highly-susceptible cultivar. The lowest mean AUDPC were for families 19-1169 and 19-1154, which were derived from a selection of ‘Otto II’, a known late-flowering cultivar. AUDPC was positively correlated with specific petiole area (r = 0.43) and inversely with days to flower (r = −0.54), green leaf index (r = −0.55), MNLI (r = −0.48) (Fig. 5D), and biomass yield (r = −0.38). Based on these associations, prolonged exposure of early flowering individuals to PM led to the loss of photosynthetic capacity and early desiccation of leaves and flowers, likely reducing biomass accumulation. Further, these results suggest that resistance to PM is quantitative and may be confounded by floral phenology. While later flowering individuals were less susceptible, the late flowering phenotype is generally not leveraged in the northeastern US because of seasonal constraints at harvest. In addition, inflorescences may not fully mature before seasonally cold temperatures, which invariably leads to yield loss. Therefore, durable genetic resistance to PM must first be identified and introgressed into elite germplasm to reduce grower inputs and maximize yield and quality traits. Selection for pest and disease resistance is still in its infancy but will likely be a key component in hemp breeding programs.

### Variation in cannabinoid profiles

Considering that all individuals in this study were derived from a common parent, cannabinoid profiles were diverse and varied substantially within and among families (Table 2; Table S3). Cannabinoid profile and ratios largely followed chemotype described in Toth *et al*. (2020), but in some families, there were significant deviations in the predicted proportions of minor cannabinoids, such as CBC (range: 0.05 to 0.1) and CBDV (range: 0.2 to 0.4). Genetic selection for novel cannabinoid profiles will require further research using segregating populations in order to identify linked or causal genetic factors (e.g., synthase copy number variation or allele-specific expression), which will aid marker development and marker-assisted selection efforts.

Of all morphological, physiological, and pathological traits measured, few were associated with cannabinoid profile (Fig. S7; Fig. S8). The strongest associations were with cannabinoid profile or chemotype (de Meijer *et al*., 2003) and foliar traits. The seed parent, ‘R4’, part of the lineage of five families, has dark, rugose, deeply serrated broad leaves and is the source of most B_T_ alleles in the trial (Table S3). In a greenhouse study, Jin *et al*. (2021) reported numerous morphological traits correlated with chemotype, however, of the chemotype I and II cultivars assayed, all were from the supposed “indica” (broadleaf) group. Conversely, Vergara *et al*. (2021) found that taxonomic designation based on leaf morphology in a segregating fiber-type (narrowleaf “sativa”) × drug-type (broadleaf “indica”) population did not correspond to cannabinoid profile, which is corroborated here. Unremarkably, significant correlations of morphological traits with chemotype reported both here and in Jin *et al*. (2021) were simply a statistical artifact – a result of population dynamics, rather than true biological relevance.

### Floral biomass yield can be predicted with few key measurements

There are key phenotypes that can be measured early in hemp development that explain a sizeable portion of the variation in biomass yield. From multiple regression, just the three variables: stem diameter, kite area, and height accounted for 67% of the explainable variation in floral biomass yield, of which 87.4% could be fully accounted for by informative predictors (Table S4). Foliar, phenology, physiology, and PM traits explained far less of the variation in biomass yield compared to stem growth and architecture, but the contribution of petiole and leaf traits to biomass yield was still significant.

The strong linear relationship of wet to sampled dry biomass (R^2^ = 0.96) in this study allowed for the accurate prediction of the former. On average, dry biomass was 30% of wet biomass and dry stripped floral biomass was 60% of dry biomass, such that ideally, at harvest, 1 kg wet biomass should yield approximately 0.18 kg dry stripped floral biomass (Table 2). Total cannabinoid yield (dry stripped floral biomass × total cannabinoid proportion) is the primary economic concern of both growers and processors. We identified four families with mean total cannabinoid yields that were significantly greater than that of the common parent: 19-1178, 19-1166, 19-1177, and 19-1162, all exceeding 150 g plant^-1^ (Fig. 6A; Table S3). The seed parents of these four high-performing families were from diversity groups 1 (T1/R4) or 2 (Cherry). It should be noted that the estimates reported for total cannabinoid yield are somewhat inflated because tissue samples from shoot apices for cannabinoid content results in an overall greater dry weight proportion compared to a random sample taken from whole plant floral biomass.

**Fig. 6.**
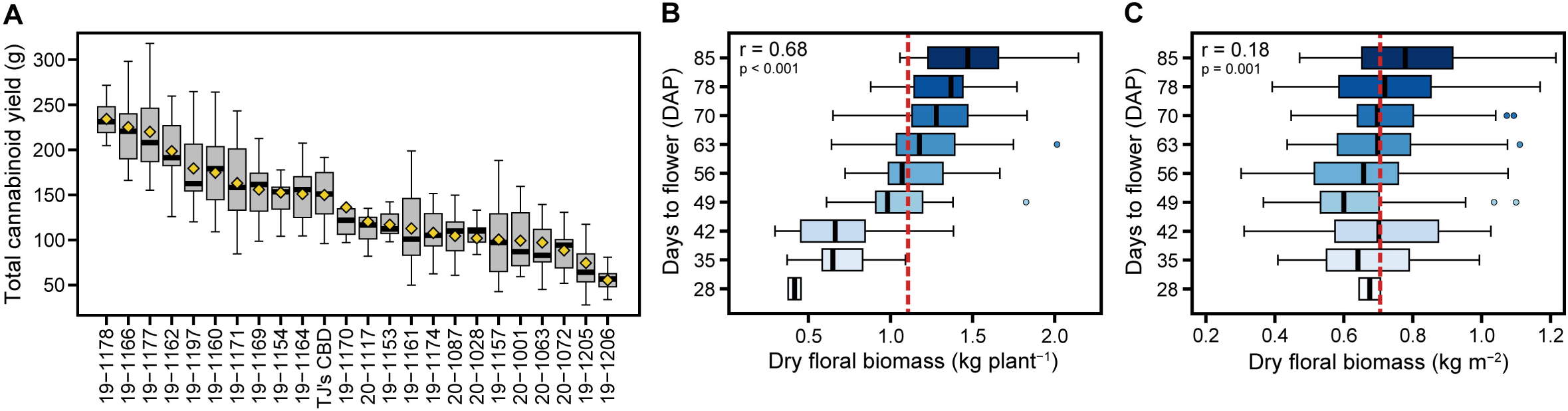
Dry floral biomass yield. Total cannabinoid yield **(A)** (dry stripped biomass × total cannabinoid proportion) by family, ordered by means (yellow diamonds). Boxplots of **(B)** dry floral biomass (kg plant^-1^) and **(C)** dry floral biomass (kg m^2^) by flowering day (DAP). Vertical dotted red lines are population means.

Compared to large, high-yielding plants, smaller plants with a greater dry stripped floral biomass to wet biomass ratio could be planted at higher densities, making harvest and processing more practical on a larger scale. Although the S_2_ families had the lowest mean biomass yield and kite area, they also had the top mean dry stripped to wet biomass ratio of 0.21, whereas the mean ratio of all other families was 0.18. Conversely, family 19-1174 was both the largest in size and highest yielding, but had a mean dry stripped to wet biomass ratio of 0.16. This ratio is inversely correlated with kite area (r = −0.72), height (r = −0.70), and growth rate (r = −0.67), and exemplifies agronomic considerations that still need to be met. Questions concerning the harvest index of hemp and its theoretical maximum are of obvious agronomic importance. Maximizing dry stripped floral biomass yield per unit area (kg m^-2^) does not necessarily favor small, early flowering cultivars, even though their dry stripped to wet biomass ratio is superior to those flowering later. In fact, when dry floral biomass is considered on the basis of yield per unit area, the correlation of days to flower and yield is virtually negligible (r = 0.18) (Fig. 6C) compared to yield per plant (r = 0.68) (Fig. 6B). Simply put, earlier flowering plants are invariably smaller but are not necessarily less space efficient. However, if early or day neutral plants are direct-seeded, they may not outcompete weeds because early flowering results in diminished growth rate and shorter height. We propose that fast growing columnar plants represent an ideotype closer to the theoretical maximum. Importantly, the use of a cultivar-specific architectural model outlined here can inform effective field design and planting density to maximize final biomass yield of hemp and will be a valuable tool for breeders and growers alike.

## Abbreviations

AUDPC: Area under the disease progress curve
CBC: Cannabichromene
CBD: Cannabidiol
CBDV: Cannabidivarin
CBG: Cannabigerol
CBL: Cannabicyclol
CBN: Cannabinol
DAP: Days after planting
DBM: Total dry biomass
DSBM: Total dry stripped floral biomass
EVI: Enhanced vegetation index
GCI: Green chlorophyll index
GDVI: Generalized difference vegetation index
HTP: High-throughput phenotyping
MNLI: Modified nonlinear vegetation index
MSAVI2: Modified secondary soil-adjusted vegetation index
NDVI: Normalized difference vegetation index
OSAVI: Optimized soil-adjusted vegetation index
PM: Hemp powdery mildew
THC: Tetrahydrocannabinol
THCV: Tetrahydrocannabivarin
UAS: Unmanned aerial system
WBM: Total wet biomass

## Supplementary Data

Supplementary data are available at *JXB* online.

Supplementary Tables S1-S4 can be found in “Carlson et al_Supplementary Tables.xlsx”.

Supplementary Figures S1-S9 in can be found in “Carlson et al_Supplementary Figures.pdf”.

***Table S1*.** Genotyping-by-sequencing sample metadata and cluster membership probabilities.

***Table S2*.** Formulas for architectural traits.

***Table S3*.** Family means, Tukey HSD groupings, and common parent deviations for all traits.

***Table S4*.** Analysis of variance from stepwise selection and relative importance of predictors for final biomass yield.

***Fig. S1*.** Genetic diversity of hemp cultivars, crosses, and U.S. feral accessions.

***Fig. S2*.** Field design.

***Fig. S3*.** Diagram of architectural traits within 2-D kite.

***Fig. S4*.** Example of calculation of mean GLI by masking operations.

***Fig. S5*.** Data processing pipeline for the extraction of phenotypic traits from RGB and multispectral data.

***Fig. S6*.** Pairwise correlations of field collected traits with aerial morphological indices on a plot-level and family-level basis.

***Fig. S7*.** Pairwise correlations of field collected traits.

***Fig. S8*.** Pairwise correlations of cannabinoid profiles and aerial indices over time.

***Dataset S1.*** Cannabinoid dataset. ***Dataset S2.*** Morphological dataset. ***Dataset S3.*** THCAS dataset.

***Dataset S4.*** UAS dataset.

***Dataset S5.*** Phylogenetic dataset.

## Acknowledgements

We thank Dr. Bill Miller (Cornell University) for assistance with formulation of silver thiosulfate, as well as Adam Berk (Stem Holdings Agri), Dr. C.J. Schwartz (Sunrise Genetics, LLC), Joseph Calderone (Emplantx, Inc.), and Edgar Winter (WinterFox Farm) for generously providing germplasm. We also thank Lauren Carlson, Allison DeSario, Deanna Gentner, Savanna Shelnutt, and Teagan Zingg, for their technical assistance, and Dr. Eric Fabio for piloting the drone.

This work was supported by the New York State Department of Agriculture and Markets through a grant (AC477) from Empire State Development, by a Federal Capacity Funds grant from United States Department of Agriculture National Institute for Food and Agriculture, and by a sponsored research agreement with Pyxus International. Bircan Taskiran was supported by a TUBITAK Fellowship.

## Author Contributions

CHC devised the experiment, collected data, performed statistical analysis, and wrote the manuscript, GMS, YJ, BT, ARC, JT, and GP contributed to data collection. CDS, JKCR, and LBS secured research funding, provided overall project design, and managed research personnel and project execution.

## Data Availability

The data that support the findings of this study are available within the paper and supplementary materials published online, and openly available in GitHub at: https://www.github.com/cornellhemp/Carlson_2021_Morphometrics/

